# SARS-CoV-2 Isolation and Propagation from Turkish COVID-19 patients

**DOI:** 10.1101/2020.04.23.056309

**Authors:** Cihan Tastan, Bulut Yurtsever, Gozde Sir, Derya Dilek Kancagi, Sevda Demir, Selen Abanuz, Utku Seyis, Mulazim Yildirim, Recai Kuzay, Omer Elibol, Serap Arbak, Merve Acikel Elmas, Selcuk Birdogan, Eray Sahin, Orhan Ozcan, Ugur Sezerman, Ercument Ovali

## Abstract

The novel coronavirus pneumonia, which was named later as Coronavirus Disease 2019 (COVID-19), is caused by the Severe Acute Respiratory Syndrome Coronavirus 2, namely SARS-CoV-2. It is a positive-strand RNA virus that is the seventh coronavirus known to infect humans. The COVID-19 outbreak presents enormous challenges for global health behind the pandemic outbreak. The first diagnosed patient in Turkey has been reported by the Republic of Turkey Ministry of Health on March 11, 2020. Today, over ninety thousand cases in Turkey, and two million cases around the world have been declared. Due to the urgent need for vaccine and anti-viral drug, isolation of the virus is crucial. Here, we report one of the first isolation and characterization studies of SARS-CoV-2 from nasopharyngeal and oropharyngeal specimens of diagnosed patients in Turkey. This study provides an isolation and replication methodology, and cell culture tropism of the virus that will be available to the research communities.

**Article Summary:** Scientists have isolated virus from Turkish COVID-19 patients. The isolation, propagation, and plaque and immune response assays of the virus described here will serve in following drug discovery and vaccine testing.

## Introduction

New coronaviruses are likely to occur periodically in humans, considering the high prevalence and wide distribution of coronaviruses, the large genetic diversity and frequent recombination of genomes, common interspecific infections, and rare outbreaks.^1,2^ The novel coronavirus pneumonia, which was named later as Coronavirus Disease 2019 (COVID-19), is caused by the Severe Acute Respiratory Syndrome Coronavirus 2, namely SARS-CoV-2^3^. It is a positive-strand RNA virus (family: Coronaviridae), showing high homology with SARS-CoV and bat coronavirus^4,5^. SARS-CoV-2 is the seventh coronavirus known to infect humans; SARS-CoV, MERS-CoV, and SARS-CoV-2 can cause severe disease, whereas HKU1, NL63, OC43, and 229E are associated with mild symptoms^6^. The novel SARS-CoV-2 is the virus behind the pandemic outbreak originating from China^7^ and this recent coronavirus outbreak (COVID-19) presents enormous challenges for global health. The first diagnosed patient in Turkey was announced by the Ministry of Health on March 11, 2020. Since then, more than ninety thousand diagnosed patients have been reported, showing the urgent need for vaccine studies and anti-viral drug discoveries. Thus, it is important to isolate and propagate SARS-CoV-2 strains from Turkish COVID-19 patients. We isolated the virus from nasopharyngeal and oropharyngeal specimens. We characterized replication properties and cell culture tropism of SARS-CoV-2 in different cell lines including green monkey kidney cell line (Vero) and Madin-Darby bovine kidney cell line (MDBK). This study provides isolation and characterization methodology, which enable research communities to propagate the isolated virus in high-titer for vaccine development and anti-viral drug screening.

## Methods

### 1. Collection and transportation of specimen

Samples were collected from the nasopharyngeal and oropharyngeal cavity of COVID-19 positive diagnosed patients according to their Real-Time PCR analysis in Acibadem Altunizade Hospital, Istanbul. Swabs were put into the transportation medium and transfered at 4°C to Acibadem Labcell Cellular Therapy Laboratory BSL-3 Units on the same day for analysis and propagation. Transfer medium contains DMEM High glucose (Thermo Fisher Scientific, USA), 2% Penicillin-Streptomycin solution (Biological Industries, Israel), and 5 µg/mL Amphotericin (Bristol Myers Squibb, USA).

### 2. Virus propagation

The propagation process was started with the 96-well plate with a Vero cell line (CCl-81, ATCC) because of the low virus titer. Firstly, Vero cells were trypsinized, centrifuged and suspended in virus media that is composed of DMEM high glucose (Thermo Fisher Scientific, USA), with 2% fetal bovine serum (Thermo Fisher Scientific, USA) and 1% Penicillin-Streptomycin-Amphotericin (PSA) solution (Pan Biotech, Germany). Cells were seeded 2.5×10^4^ into each well. Then, 100 µl of virus sample was added to the 1^st^ line, and 50 µl of the virus media added to other wells of plate. Serial dilution was done starting from the 1^st^ line to 12^th^ with 50 µl of virus solution and incubated at 37°C. Each day cytopathic effect was recorded under inverted microscope (Leica Microsystems) and after its detection; cells were scraped from the well and transferred to the 24-well plate with the Vero cell line in 25×10^4^ concentrations per well. The cytopathic effect was observed and cells transferred to the T75 tissue culture flask. Before adding the virus inoculation, Vero cells were seeded in T75 tissue culture flask in 5×10^6^ concentration and virus inoculum were transferred to monolayer cells. At this point, the average number of days for cytopathic effect can be known as 3-4 days. For higher propagation, virus inoculum was transferred to the T300 tissue culture flask.

### 3. Isolation of Viruses

Infected cells were observed each day and the following analysis was performed to confirm SARS-CoV-2.

#### Real-Time PCR

Total RNA isolations were carried using Direct-zol RNA Miniprep Kits (Zymo Research, USA), and concentrations were determined using Qubit fluorometer with the Qubit RNA HS Assay (Thermo Fisher Scientific, USA). cDNA synthesis was performed using random oligonucleotides. Two LAMP assay reactions were set-up in CFX-96 (Bio-Rad, USA) for each sample with SARS-CoV-2 LAMP primer mix and internal control LAMP primer mix. EvaGreen fluorescent dye (Biotium, USA) was added to each reaction mixture for real-time LAMP detection. Melt curve analysis and %2 agarose gel electrophoresis were used for the evaluation of the results. Here, the virus N gene was used as a target gene in SARS-CoV-2 and actin as an internal control.

#### Transmission Electron Microscopy (TEM)

Infected Vero cells were scraped from the flask and centrifuged at 300xg for 10 minutes. The cell pellet was rinsed with 0.1M phosphate buffer saline (Thermo Fisher Scientific, USA) centrifuged at 300xg for 10 minutes. Pellet was fixed 2.5% glutaraldehyde in 0.1 M Phosphate Buffer Saline (PBS) (pH 7.4) for at least 2 hours at room temperature. The fixed block was washed in 0.1 M phosphate buffer three times for 15 minutes. Post fixation was done in 1% osmium tetroxide in 0.1 M phosphate buffer 1-2 hours at room temperature. In this step, 2% aqueous OsO4 and 0.2M phosphate buffer were mixed in equal volume and used immediately. En Bloc Staining was done to enhance contrast. Bloc was washed at least 3-5 times of 10 minutes each in distilled water to remove all excess phosphate ions, therefore, preventing uranyl acetate (UA) from being precipitated. En bloc was stained with 2% aqueous uranyl acetate for 1 hour at room temperature or 2 hours at 4 °C in dark to prevent uranyl acetate from being precipitated as UA is photo reductive. It was washed 2-3 times of 5 minutes in distilled water. In the dehydration process, bloc was put in 50% ethanol for 10-15 min. Then, it was put in 70% ethanol for 10-15 min. If it is necessary, the bloc may be stored overnight at this stage. The alcohol ratio was increased to 95% and put in it for 10-15 min then transferred into 100% ethanol 3 times for 15 min. Bloc was put 100% Propylene oxide 3 times for 15 min. The sample was transferred in 1:1 EMBed 812 and Propylene Oxide overnight at room temperature in tightly capped vials to prevent moisture from coming into the specimen. EMBed 812 (100%) was straight for 1-2 hours at room temperature. The caps were removed from vials to allow any remaining propylene oxide to evaporate. The sample was labeled and embedded in beam capsules or embedding molds. It was polymerized in 60-70 °C (∼65 °C) oven for 24-48 hours. The sample was viewed in different scales.

#### Plaque Assay

Plaque test was performed based on general procedure with small modifications as following. Vero cells were seeded for 6-well plate (1×10^5^cells/well) and leaved for 24 h incubation at 37°C. After cell incubation period, virus titration was prepared in Log10 serial dilutions. Cells were washed with cell media (DMEM High, Thermo Fisher Scientific, USA) that contain only 1% PSA (Pan Biotech Germany). Virus titrations were inoculated to cells in 1 ml volume, leaved for the 1-hour incubation, and plates were shaken gently in 20 minutes. In this period also plaque medium that contains 2% agarose and media (2% FBS and 1% PSA) in 1:1 volume ratio were prepared and put into the 56°C water bath for 30 minutes. After virus inoculation, excess viruses were removed and 1 ml plaque medium added on infected cells. The plaque medium should warm avoid the damage of the cell monolayer. Plates were left under the hood for 15 minutes without a cap and then put into 37 °C incubator for 7-10 days. For the counting of the plaque firstly cells were fixated with 10% of paraformaldehyde for 1 hour. Agarose layers were removed from the well and then 500 µl of crystal violet solution was added to each well and incubated on a shaker for 5-10 minutes. Crystal violet was discarded and plates were washed with tap water without disrupting cell monolayer until the water was clear. Plates were left on a paper towel till dry. Plaques were imaged under an inverted microscope.

### 4. Immunological Response

#### Inactivation of Virus

Infected Vero cells were scraped from the flask and centrifuged at 1000 rpm for 10 min. The cell pellet was rinsed with 0.1M phosphate buffer saline (Thermo Fisher Scientific, USA) centrifuged at 1000 rpm for 10 min. Pellet was fixed 2.5% glutaraldehyde (GA) in 0.1 M Phosphate Buffer Saline (PBS) (pH 7.4) for at least 2 hours at room temperature. Virus solution (15 ml) was added to Amicon Ultra 50,000 KDa NMWL and centrifuge at 4,000xg for 15 minutes to wash from excess GA. For washing sample was diluted with 1ml of PBS and centrifuge for 15 minutes at 4,000xg. The sample was diluted with 1ml of PBS and centrifuge for 3 minutes at 4,000xg. The concentrator was transferred to a clean tube and centrifuge at 1,000xg for 2 minutes to collect the concentrated sample. The sample was diluted in serum physiologic.

#### PBMC Culturing

Three healthy adult bloods were obtained from Acibadem Labcell Cellular Therapy Laboratory. Following the isolation of peripheral mononuclear cells (PBMC) by overlaying blood on Ficoll-paque plus (GE Healthcare), the serially diluted GA-inactivated SARS-CoV-2 samples were incubated with the PBMCs for 48hr in T cell medium (6% Human AB Serum and 1% Pen/Strep, Texmacs medium). Immunocult human CD3/CD28 T cell activator (STEMCELL Technologies, USA) was used to stimulate the PBMCs as a positive control for immunological activation.

#### Flow cytometry

Immune cell subtypes and their activation levels were determined by Miltenyi MACSQuant flow cytometry analysis using αCD3-PE, αCD19-PE.cy7, αCD56-FITC, αCD4-Viogreen, αCD8-Vioblue, αCD25-APC and αCD107a-PE.cy5.5 (Miltenyi).

#### Interferon-gamma ELISA

Three human PBMC’s were added to different inactivated SARS-CoV-2 virus diluted to five different dilutions. After 48 hours, supernatants were collected and cytokine levels assed with Human IFNγ ELISA Kit (ThermoFisher Scientific, Vienna, Austria) according to the manufacturer’s instructions. The supernatants (50 µl) were added into 450 µl sample diluents (TexMACs) and 50µl test samples, positive control (immunocult-activated) and negative control were added into wells. ELISA plate was measured at 450 nm, and 550 nm using a Microplate reader (Bmg Labtech, Germany).

## Results

### Propagation and Genetic Confirmation of SARS-CoV-2 collected from COVID-19 patient

The SARS-CoV-2 including swab specimens collected from COVID-19 diagnosed patients were quickly transferred in the same day to the laboratory and incubated with Vero cell in 96-well plate as mentioned in the method. Because of low titer viruses, initial small size culture led easy recovery. Upon observation of the first cytopathic effect, supernatant and the cells were transferred to a 24-well plate with suspension form of Vero cell. Through one week, propagation of the virus was followed and increasing cytopathic effects were recorded (**Fig. 1A**). As Bovine Coronavirus (B-CoV) is incubated with a bovine kidney cell line (MDBK)^8^, we wanted to test propagation of the SARS-CoV-2 with this cell line. It showed cytopathic effect faster than Vero cell line (**Fig. 1B**). Next, we wanted to identify the propagated virus as SARS-CoV-2 using RT-PCR along with SARS-CoV-2 N gene specific primers. Results shown in **Fig. 1C** and **1D** proved the presence of SARS-CoV-2 in the propagated samples.

**Figure 1:**
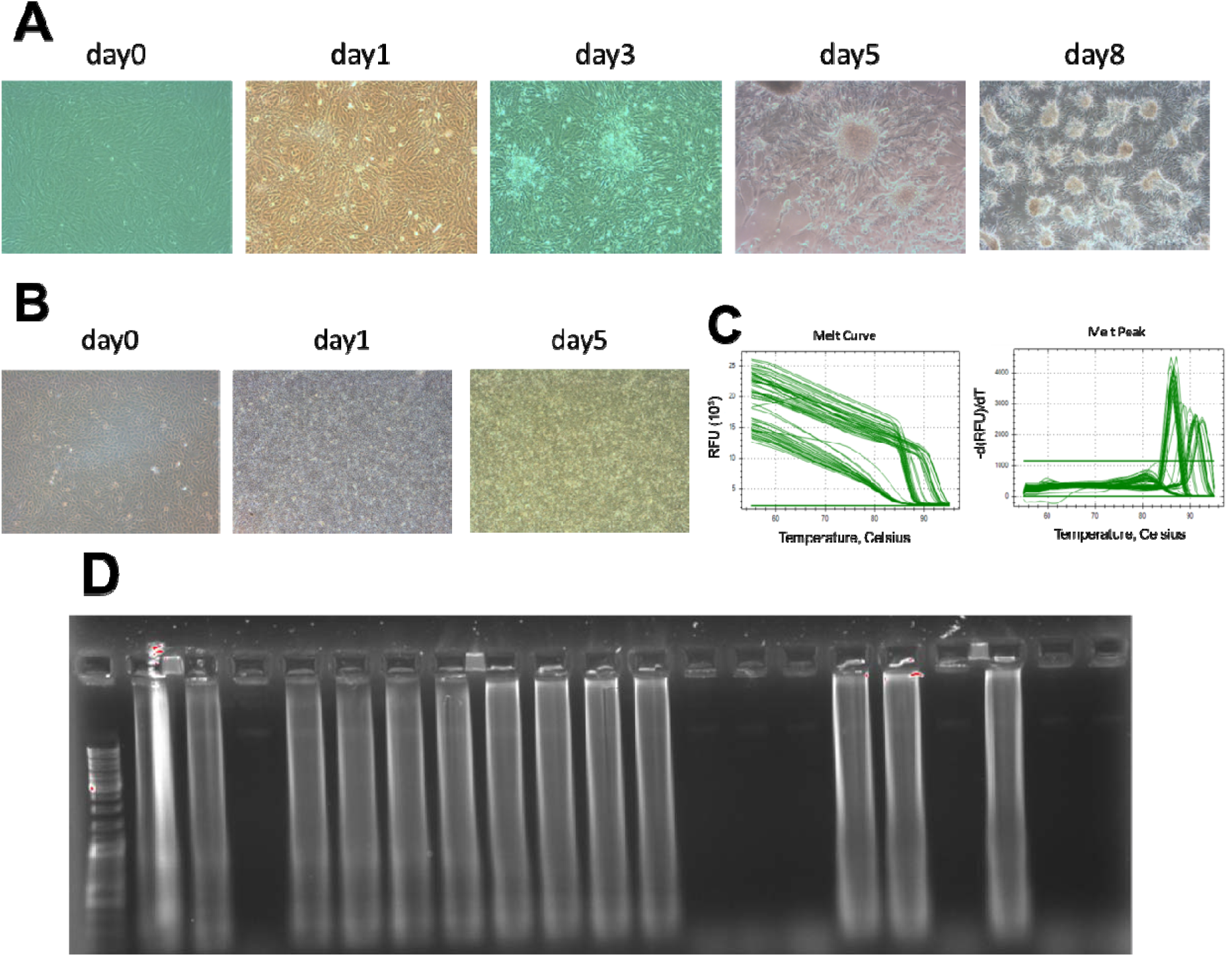
SARS-CoV-2 Propagation and Genetic Confirmation. **A**. Images of virus propagation in Vero cell line. **B**. Images of virus propagation in MDBK cell line. **C**. Graphics of amplification curve (left) and melt peak (right) of SARS-CoV-2 N gene and β-actin control. **D**. Agarose gel electrophoresis of the LAMP products (13 out of 18 collected propagated samples identified as SARS-CoV-2 positive) and the products resulted from cut of the LAMP products with AccI restriction endonucleases separated on 2% agarose gel and stained with ethidium bromide.

### Transmission Electron Microscopy imaging and Plaque assay of SARS-CoV-2

Afterwards, a detailed scanning was performed with TEM analysis to show existence of the SARS-CoV-2 (**Fig. 2**). We could easily detect spike protein of SARS-CoV-2 in the images, convincing us the virus propagated in the cell line (**Fig. 2**). Plaque assay is another way to show virus infectivity^9^. Therefore this assay was performed on Vero and MDBK cell lines for isolated and propagated virus. Although the literature^10^ suggests that plaque formation were determined in 72hr, the plaques were observed in our experiments after 6 days in both cell lines (**Fig. 3**). The results confirmed our methodological approach for isolation and propagation of SARS-CoV-2.

**Figure 2:**
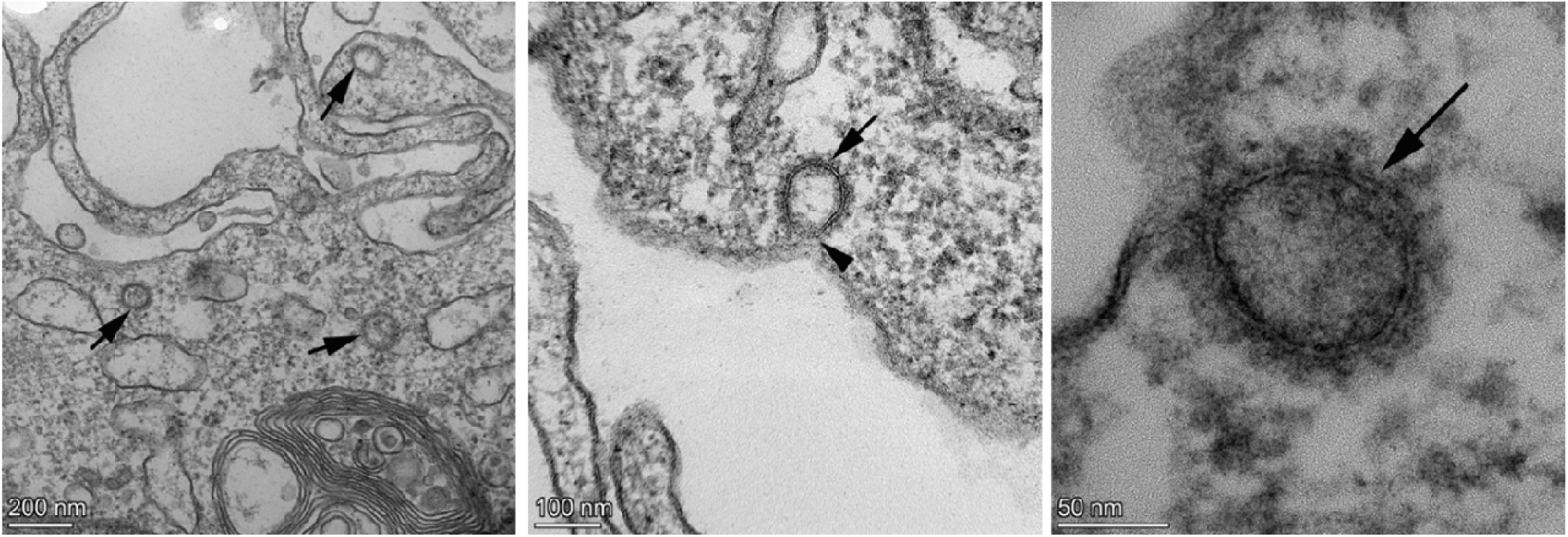
Transmission electron microscopy images of SARS-CoV-2 propagation in Vero cell line. Arrows show the virus in the images taken at different scales (200-100-50 nm).

**Figure 3:**
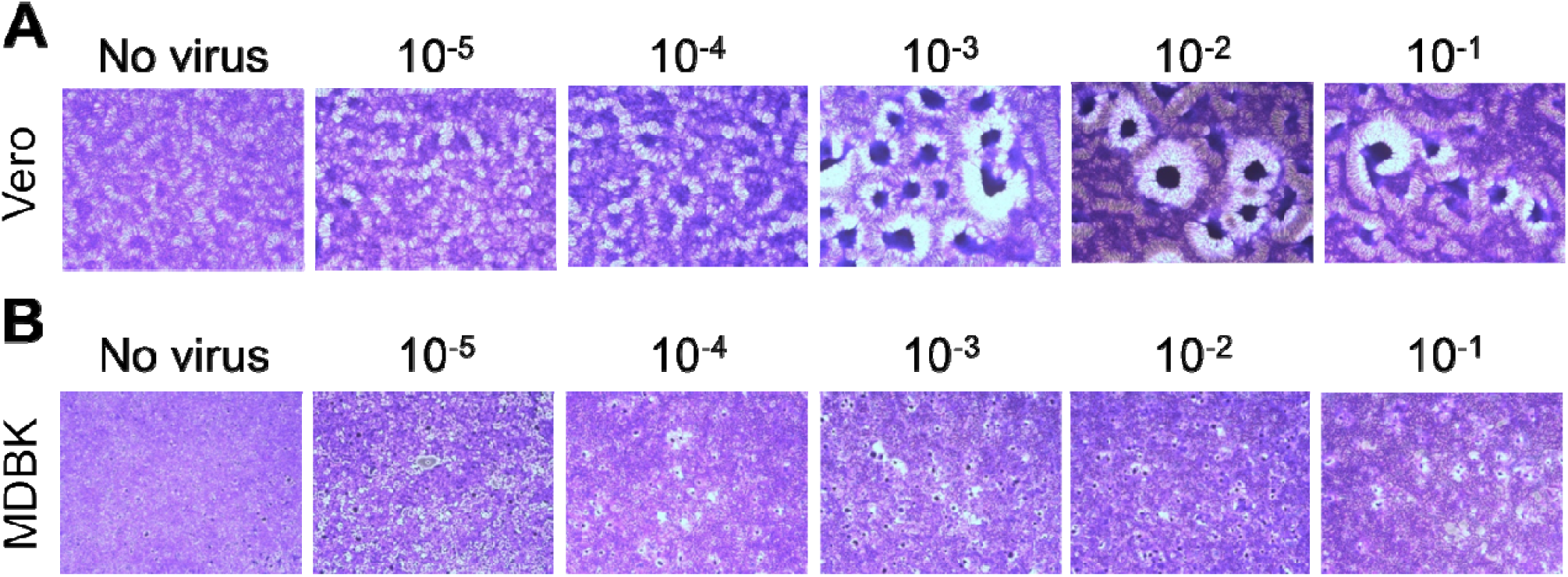
Plaque assay of SARS-CoV-2 in A. Vero and B. MDBK cell lines with different virus titers. Images are taken in 6^th^ day of the assays.

### Immunological Response of Inactivated Virus

PBMC from healthy donors are a useful tool for assessment of host response upon pathogen infection. The cells can be stimulated by encountering SARS-CoV and recruit immune cell subtypes such as natural killer T (NKT), T, and B cells to the site of inflammation to give early adaptive immune response^11^. Therefore, we performed immunological assays to show impact of GA-inactivated SARS-CoV-2 on frequencies and activation capacities of the cell subtypes. We incubated healthy adult PBMCs with the inactivated virus in a dose-dependent manner for 48hr and assessed their proportions and expression level of activation markers (CD25 and CD107a) using flow cytometry as shown in **Fig. 4A**. We determined an increase in frequency of CD3+ T cells, especially CD3+ CD4+ T helper cell, except CD3+ CD8+ cytotoxic T cells at the highest concentration of the virus (**Fig. 4B-C)**. On the other hand, CD19+ B cell proportion has decreased upon increasing of the virus concentration, suggesting activation of the immune cells (**Fig. 4B-C)**. Afterwards, we assessed activation of the T cells and NKT cells, and we determined a significant upregulation of CD25 on CD3+ CD56+ NK T cells and an increase of CD25 level in CD3+ CD4+ T helper cell but not in CD3+ CD8+ cytotoxic T cells (**Fig. 4B-C)**. However, we could not detect expression of a degranulation marker (CD107a) on the cells (data not shown). These results led us to determine another immunological stimulus marker (IFNγ secretion) using collected supernatant of the samples after 48hr. We determined a significant increase in IFNγ secretion (**Fig. 5**) using ELISA, which confirm occurrence of the immune response with chemically inactivated SARS-CoV-2.

**Figure 4:**
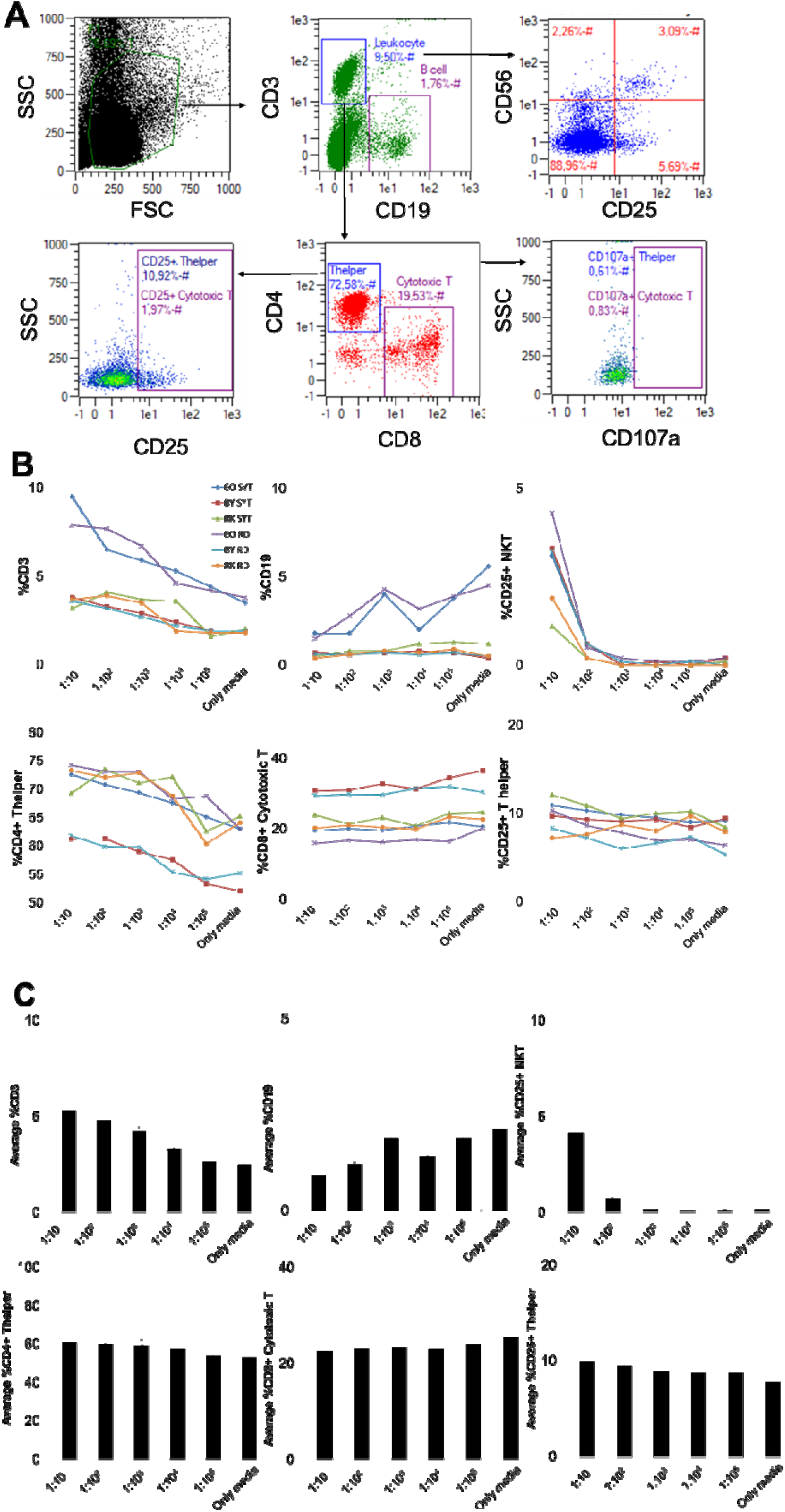
Stimulated Immune response by GA-inactivated SARS-CoV-2. **A**. Flow cytometry plots that show immune cell subtypes (CD19+ B, CD3+ CD4+ T helper, CD3+ CD8+ cytotoxic T, and CD3+ CD56+ Natural Killer T (NKT) cells) and their activation (with αCD25 and αCD107a). Graphs that show **B**. frequencies (linear graphs) and **C**. averages (bar graphs) of the immune cell subtypes and their activation from three healthy donor peripheral blood mononuclear cells incubated with two different SARS-CoV-2 samples.

**Figure 5:**
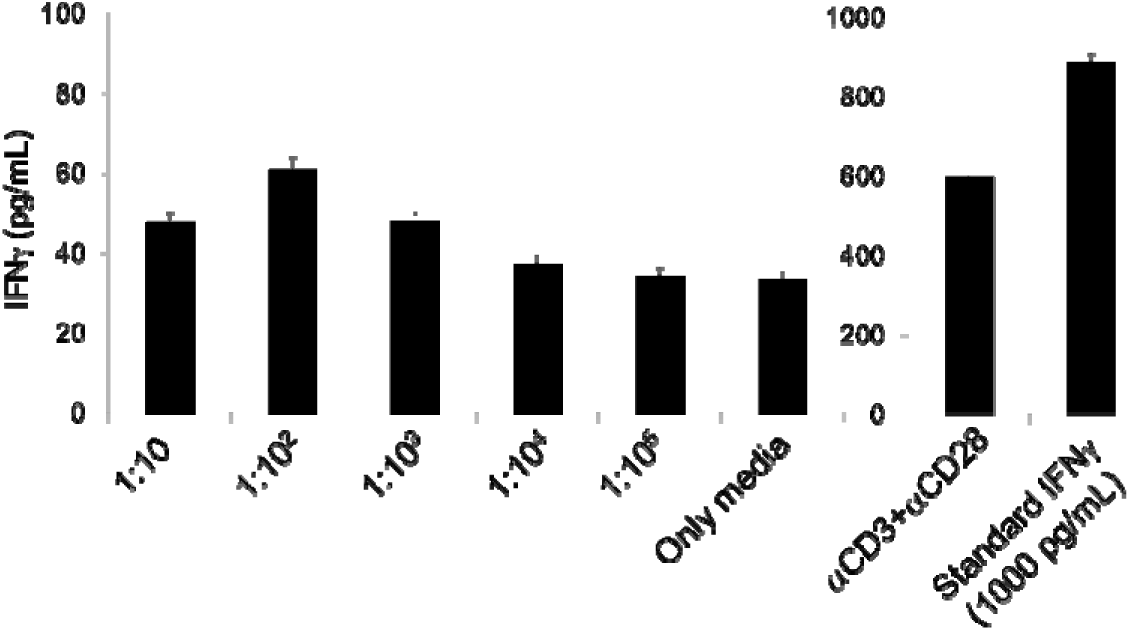
IFNγ secretion capacity of the stimulated PBMCs by SARS-CoV-2 in a dose-dependent manner. The graphs show IFNγ secretion (pg/mL) upon 48hr incubation of the PBMCs with the virus (left graph) or with αCD3+αCD28 Immunocult (right graph).

## Discussion

Coronaviruses are enveloped RNA viruses that are widely distributed among humans, other mammals, birds, which cause respiratory, enteric, hepatic and neurological diseases^12,13^. After the first outbreaks of unexplained pneumonia in Wuhan, China, a new coronavirus was detected as a disease-causing agent in late 2019 and January 2020^14^. In April, 2020, above two million cases have been reported from 26 countries, including China^15^. The emergence of coronaviruses at regular intervals poses an important threat to human health and economy of countries. Ironically, even after a decade of research on the coronavirus, there are no licensed vaccines or therapeutic agents to treat coronavirus infection. This emphasizes the urgent need to develop effective vaccines to prevent future outbreaks^16^ and therefore isolation of the SARS-CoV-2 viruses become important issues.

In the process of specimen collection from patient and transfer to the laboratory, the process should be started and virus cultured immediately because lots of virus became inactivated. Therefore, we tested several transfer solutions to safe infectious SARS-CoV-2 during the transportation and we decided FBS-free media is the best for protecting the infectivity of the virus. Also, before culturing the cells with the specimens, we wash the cells with FBS free media to remove excess of fetal bovine serum. Then, to propagate virus, cells were resuspended with a virus media including 2% FBS, which enabled much easier propagation of the virus than the media including 10% FBS. Afterwards, we tested Vero and MDBK cell lines to observe cytopathic effects and it was determined that the spreading of SARS-CoV-2 occurred in MDBK cells faster than in Vero cells. Also, the cytopathic effect in Vero cell line was observed as clump form while in MDBK cell line as a single cell bases. Plaque assay is a reliable way to show an infectious form of viruses. Therefore, in this study, we performed plaque assay to see infectivity of SARS-CoV-2. During the plaque assay processes, we observed that monolayer form of Vero cells behave more sensitive than that of MDBK cells, which makes difficulties in setup of the assay. Therefore, MDBK cell line is suggested for efficient plaque assay for virus titration and antiviral drug screening. Furthermore, studies reported that plaque formation time for SARS-CoV-2 was 72h, however; in our study, plaque formation was observed in 6^th^ day of the infection, probably because of the virus low titer. Thus, we suggest waiting for 10 days post infection to determine titer of the virus by counting plaques.

Peripheral blood mononuclear cells are used to test immune response stimulated by pathogens. To determine immune cell activation through the proteins of inactivated SARS-CoV-2, we incubated PBMCs with chemically-inactivated virus. The study shows us that T cells and NKT cells can be stimulated with the virus and can be assessed significantly in *in vitro* setup. We also observed a decrease in B cell population in higher concentrations of the inoculums; probably activated and proliferated T cells interfere with B cell proliferation^17^. To further confirm the activation, we determined IFNγ secretion from the stimulated cells in PBMCs. These preliminary results suggest that early immune response tests can be performed *in vitro* in the process of vaccine and anti-viral drug developments.

To conclude, this study can be referenced for researchers who aimed to isolate and propagate SARS-CoV-2 to work in molecular biology studies including genomics, proteomics, and gene editing like CRISPR, antisense peptide and small interference RNA (siRNA) therapies.

## Acknowledgement/Disclaimers/Conflict of interest

None

## Acknowledgement

The authors would like to thank Prof. Fikrettin Sahin and Prof. Dilek Telci from Genetic and Bioengineering Department, Yeditepe University, Istanbul for their comments and their insightful suggestions for the study.

